# Growth allometry and dental topography in Upper Triassic conodonts support trophic differentiation and molar-like element function

**DOI:** 10.1101/2021.06.10.447946

**Authors:** Valentin Kelz, Pauline Guenser, Manuel Rigo, Emilia Jarochowska

## Abstract

Conodont elements have high rates of morphological evolution, but the drivers of this disparity are debated. Positive allometric relationships between dimensions of food-processing surfaces and entire P_1_ elements have been used in to argue that these elements performed mechanical digestion. If involved in food processing, the surface of the element should grow at a rate proportional to the increase in energy requirements of the animal. This inference of function relies on the assumption that the energy requirements of the animal grew faster (≅ mass^0.75^) than the tooth area (≅ mass^0.67^). We re-evaluate this assumption based on metabolic rates across animals, and calculate the allometry in platform-bearing P_1_ elements of Late Triassic co-occurring taxa, *Metapolygnathus communisti* and *Epigondolella rigoi*, using 3D models of ontogenetic series. Positive allometry is found in platform and element dimensions in both species, supporting a grasping-tooth hypothesis, based on the assumption that metabolic rate in conodonts scaled with body mass similarly to that in fish and ectoterms. We also calculate the curvature of P_1_ platform surface using the Dirichlet Normal Energy (DNE) as a proxy for diet.DNE values increase with body mass, supporting the assumption that conodont metabolic rates increased faster than mass^0.67^. We finally find that adults in both taxa differ in their food bases, which supports trophic diversification as an important driver of the remarkable disparity of conodont elements.

## Introduction

Conodonts are extinct, eel-shaped marine animals that lived from the mid-Cambrian to Early Jurassic (Du et al. 2020). They are early vertebrates, either stem-gnathostomes (Donoghue et al. 1998) or stem-cyclostomes (Miyashita et al. 2019), and they are distinguished by an extensive fossil record (Foote and Sepkoski 1999; Donoghue 2001a). The majority of the conodont fossil record consists of the phosphatic elements forming the feeding apparatus of the animal (Schmidt 1934; Scott 1934; Purnell et al. 2000). Conodont elements were retained throughout the life of an individual (Donoghue and Purnell 1999a), recording periodic growth through the apposition of crown tissue lamellae (Zhang et al. 1997; Dzik 2008). Soft tissues are rarely found and have, so far, not revealed a great diversity of body forms. Conodont taxonomy and functional morphology are thus based on their feeding elements (Mazza et al. 2012b for the Upper Triassic and references therein). Patterns interpreted to be produced by elements shearing on the surface and repaired during the animal’s growth suggest macrophagy (Purnell 1995; Donoghue and Purnell 1999a; Shirley et al. 2018). An active, predatory lifestyle is supported by the discovery of a conodont specimen with preserved extrinsic eye musculature (Gabbott et al. 1995), which was interpreted as indicative of conodonts having pattern vision (Purnell 1994). Calcium isotope analyses indicated that Late Devonian conodonts were first-level zooplanktivore and piscivore consumers (Balter et al. 2019), suggesting that in this period, conodonts did not live a purely predatory lifestyle. However, despite several publications related to this topic, the diet of conodonts and its evolution are far from resolved. Conodonts changed their apparatus structure and disparity across their stratigraphic range (Dzik 1991, 2015), possibly reflecting their evolving niches as marine ecosystems increased in complexity (Klug et al. 2010; Ginot and Goudemand 2019). Under the assumption that conodont element morphology is an adaptation to their diet (Jones et al. 2012a; Ginot and Goudemand 2019; Guenser et al. 2019; Petryshen et al. 2020), the disparity suggests changing trophic position throughout the existence of the lineage.

Calcium isotope analysis indicated that trophic niches overlapped, suggesting competition between some taxa (Balter et al. 2019). On the other hand, Sr/Ca ratios in Silurian conodont assemblages indicate differences between species (Terrill et al. 2022), possibly reflecting trophic niche differentiation through disruptive selection. Since there is no direct evidence of conodont food base, trophic diversity of conodonts must be inferred from proxies, e.g. by evaluating morphological and functional diversity of food-processing elements.

Here we revisit the test between two hypotheses on the function of conodont apparatus: the suspension-feeding model (Nicoll 1985, 1987) *versus* the grasping-tooth model (Aldridge et al. 1987; Purnell and von Bitter 1992). In the grasping-tooth model and its subsequent modifications, S and M elements, positioned in the anterior part of the mouth, are interpreted to perform a grasping function, whereas P elements, placed in the posterior part of the apparatus, in the pharynx of the animal, have a function similar to molars of mammals (Purnell and von Bitter 1992; Purnell 1994; Donoghue and Purnell 1999b; Goudemand et al. 2011). In the suspension feeder model by Nicoll (1987), elements would be covered in tissue and S and M elements would filter particles and create current, whereas P elements would only lightly mash food.

Several analytical methods have been proposed in favor of the grasping-tooth model: (1) observation of microwear patterns and damage of conodont elements, produced *in vivo* (Purnell 1995; Martínez-Pérez et al. 2014a; Shirley et al. 2018); (2) occlusion models (Donoghue and Purnell 1999*b*; Jones et al. 2012a; Martínez-Pérez et al. 2014b, a); (3) Finite Element Analysis (FEA) (Jones et al. 2012b; Martínez-Pérez et al. 2014a, 2016); (4) histological adaptation (Donoghue 2001b; Jones et al. 2012a); and (5) growth allometry (Purnell 1993, 1994), which is central in this study.

Allometry describes proportional relationships of body parts, usually of the size of an organ relative to the total size of the organism. Proportional growth, whereby the growth of an organ and the size of the animal increase at the same rate, is called isometry. Positive allometry then describes the organ growing at a faster rate than the rest of the animal. Negative allometry, conversely, describes the organ growing at a slower rate than the rest of the body (e.g., Gould 1966; Alberch et al. 1979; Klingenberg 1996). Methods of investigation listed in the previous paragraph have focused on elements termed P_1_. Those elements are typically the largest and the most robust in the apparatus and, thus, inferred to have the most pronounced dental function.

They also have the highest morphological evolutionary rate and, thus, present the most diverse shapes within the conodont apparatus. Among the possible P_1_ morphologies, some of them bear a platform, which is inferred to be a food-processing surface (Purnell 1995). Positive allometry of the platform area relative to the total length of the P_1_ element in Carboniferous taxa *Idiognathodus* sp. and *Gnathodus bilineatus* has been used in seminal studies by Purnell (1993, 1994) to test between the two hypotheses on feeding in conodonts. The test used by Purnell (1993, 1994) was devised based on the understanding of growth allometry at the time, which led to the following assumptions: (1) a tooth’s ability to process food is proportional to its area; (2) any tooth linear dimension increases as any other tooth linear dimension (isometric slope = 1); (3)tooth area increases as the square of its linear dimensions (isometric slope = 2) or as its volume to the power of 0.67; (4) the animal’s metabolic rate and thus energy requirements increase as its volume to the power of 0.75 (Gould 1966, 1975). Based on these assumptions, the dental function of P_1_ conodont elements, analogously to molar teeth in mammals, should manifest itself in positive allometry of the food-processing area *versus* the element’s linear dimensions to meet the food requirements (Figure 1).

**Figure 1:**
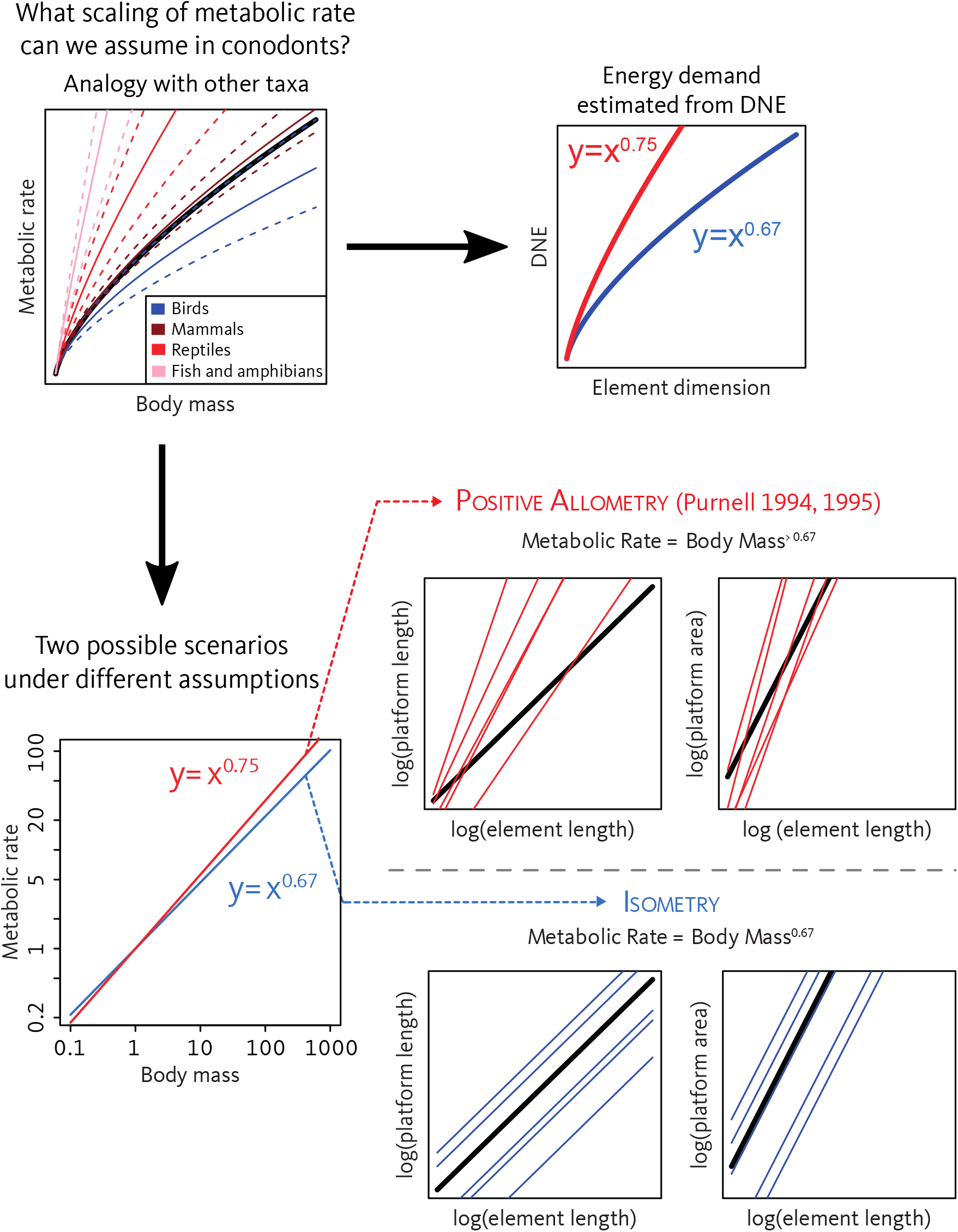
Theoretical framework of the present study. Data from birds, mammals, reptiles, fishes and amphibians were extracted from Glazier (2009); dashed lines represent 95% CI.

In the present study we first apply the same allometric evaluation of P_1_ elements as used by Purnell (1993, 1994) to two Late Triassic platform-bearing conodont species *Metapolygnathus communisti* Hayashi, 1968 (Hayashi 1968) and *Epigondolella rigoi* Kozur, 2007 (Noyan and Kozur 2007). However, we also revisit assumption 3 (previous paragraph) based on the research progress since the original studies by Purnell (1993, 1994) and examine the implications when they are violated (see Discussion). Assumption 3 is that tooth area can be approximated by a two-dimensional surface, such as when analyzing photographs of teeth in a plane view. An alternative way of increasing the surface is introducing topography – a solution found in mammals (Santana et al. 2011; Pérez-Ramos et al. 2020). Conodonts have developed intricate topographies on the platform surface of their P_1_ elements, which would not be identified as increased areas when approximating the total area with an ellipse described by the element’s linear dimensions (Purnell 1993, 1994). Therefore, here we use 3D meshes to calculate the three-dimensional platform area and thus account for the topography. To evaluate the importance of dental topography in conodonts, we calculate Dirichlet Normal Energy (DNE), which measures the curvature and morphological irregularity of a surface. Essentially, DNE measures how much a given surface differs from a plane (Bunn et al. 2011). Surface topography is an important feature of teeth because it helps break down food to satisfy the demand for energy (Bunn et al. 2011). Then, a particular DNE value of the occlusal surface of a tooth (or tooth-like buccal elements in conodonts) should reflect a particular diet. However, DNE has, until now, been only used to analyze skeletal parts of mammals, especially primate teeth (Bunn et al. 2011; Godfrey et al. 2012; Winchester et al. 2014; Prufrock et al. 2016; Berthaume and Schroer 2017; López-Torres et al. 2018; Pampush et al. 2019; Fulwood 2020; Li et al. 2020; Cuesta-Torralvo et al. 2021), but also those of marsupials (Lang et al. 2022), carnivorans (hyenas, bears) (Pérez-Ramos et al. 2020; de Vries et al. 2021; Lang et al. 2022), scandentians (tree shrews) (Selig et al. 2019), rodents (Prufrock et al. 2016; Renaud and Ledevin 2017; Vermeer 2019; de Vries et al. 2021), chiropterans (Pellegrom 2019; López-Aguirre et al. 2022; Villalobos-Chaves and Santana 2022), multituberculates (Robson 2018), artiodactyls (suids) (Rannikko et al. 2020), eulipotyphles (hedgehogs) (Vitek et al. 2021) and teeth of one mammal stem group (Harper et al. 2019). DNE is defined in such a way that it is independent of scale. Thus, any change with the linear dimensions of the animal might be interpreted as a functional change and not a scaling effect (Pampush et al., 2022). DNE values of platform surface of P_1_ elements have the potential to decipher the diet of a conodont species. Preliminary dental topographic analyses, different than DNE, have been applied to conodont elements suggested that conodonts might had different diets through their evolution (Purnell and Evans 2009; Stockey et al. 2021, 2022). These recent findings support the potential of dental topographic methods for resolving conodont diets and DNE is a novel method to test functional hypotheses.

The aims of this study are to:

1. Test the null hypothesis of isometric growth against the alternative of positive allometric growth of P_1_ elements in a new set of taxa, to evaluate whether the relationship previously observed in Carboniferous taxa by Purnell (1993, 1994) holds more widely in the class Conodonta;
2. Test the null hypothesis of an absence of relationship between DNE values of P_1_ platform surface and P_1_ element dimensions (i.e. element length, platform length and platform area). If such a relationship exists, by fitting the data with a power law, it is discussed in the light of the energy requirement of the animal;
3. Revise the assumption that the metabolic rate in conodonts increased faster with body mass than tooth area and evaluate how violation of this assumption would affect the outcome of the allometric analysis by Purnell (1993, 1994) and carried out here;
4. Test the hypothesis that differential morphologies in P_1_ elements of the co-occurring taxa *M. communisti* and *E. rigoi* reflect different food bases within the ecosystem by comparing DNE values between P_1_ platformsurfaces in adult specimens.

## Materials & Methods

### Material

We studied two growth series of the ozarkodinid conodont species *Metapolygnathus communisti* and *Epigondolella rigoi* from the Pizzo Mondello section in western Sicily, Italy. They were collected from a section of 430 m thick marine limestone dated to the upper Carnian to upper Norian (Mazza et al. 2012a). Twenty-seven P_1_ elements of *M. communisti* and 23 P_1_ elements of *E. rigoi* were used, separated into six growth stages (GS) based on the maturity of morphological characters of the platform (Mazza and Martínez-Pérez 2015). The six growth stages are GS1 – early juvenile, GS2 – juvenile, GS3 – late juvenile, GS4 – early adult, GS5 – adult and GS6 – late adult (Mazza and Martínez-Pérez 2015) (Table 1). At Pizzo Mondello, *M. communisti* occurs from the upper Carnian to the lower Norian (from c.a. -227.5 Ma to c.a. - 226.5 Ma) (Mazza et al. 2012a, 2018; Ogg et al. 2020). The specimens range from late juvenile to late adult (i.e., from GS3 to GS6), though mature elements are more abundant (Table 1). The stratigraphic range of *E. rigoi* at Pizzo Mondello is from the lower Norian to the middle Norian (from c.a. -227 Ma to c.a. -216 Ma) (Mazza et al. 2010, 2012a; Ogg et al. 2020), a longer interval than *M. communisti*. Elements range from GS2 to GS5: earlier ontogenetic stages are sparse (Table 1). These specimens have an average colour alteration index (CAI) of 1, suggesting minimal post-depositional heating (Epstein et al. 1977; Nicora et al. 2007; Mazza et al. 2012a). The studied elements are housed in the collection of the Dipartimento di Scienze della Terra “A. Desio” of the Università degli Studi di Milano. The whole conodont collection from the Pizzo Mondello section is housed in Milan and Padova (Department of Geosciences, University of Padova).

**Table 1.** Numbers of conodont P1 element specimens by growth stages used for the study. Abbreviations: GS – Growth stage.

|  | *Metapolygnathus communisti* | *Epigondolella rigoi* |
| --- | --- | --- |
| GS2 | 0 | 1 |
| GS3 | 3 | 1 |
| GS4 | 2 | 6 |
| GS5 | 7 | 15 |
| GS6 | 15 | 0 |

### Methods

#### Scanning

The specimens were scanned with a resolution of 1 μm with a microtomograph nanotomS (General Electric) of the AniRA-ImmOs platform, SFR Biosciences (UMS 3444), Ecole Normale Supérieure de Lyon, France. Amira^©^ software was used for the 3D reconstruction (Guenser et al. 2019; Figure 2). The meshes are available on MorphoBank: http://morphobank.org/permalink/?P4048

**Figure 2:**
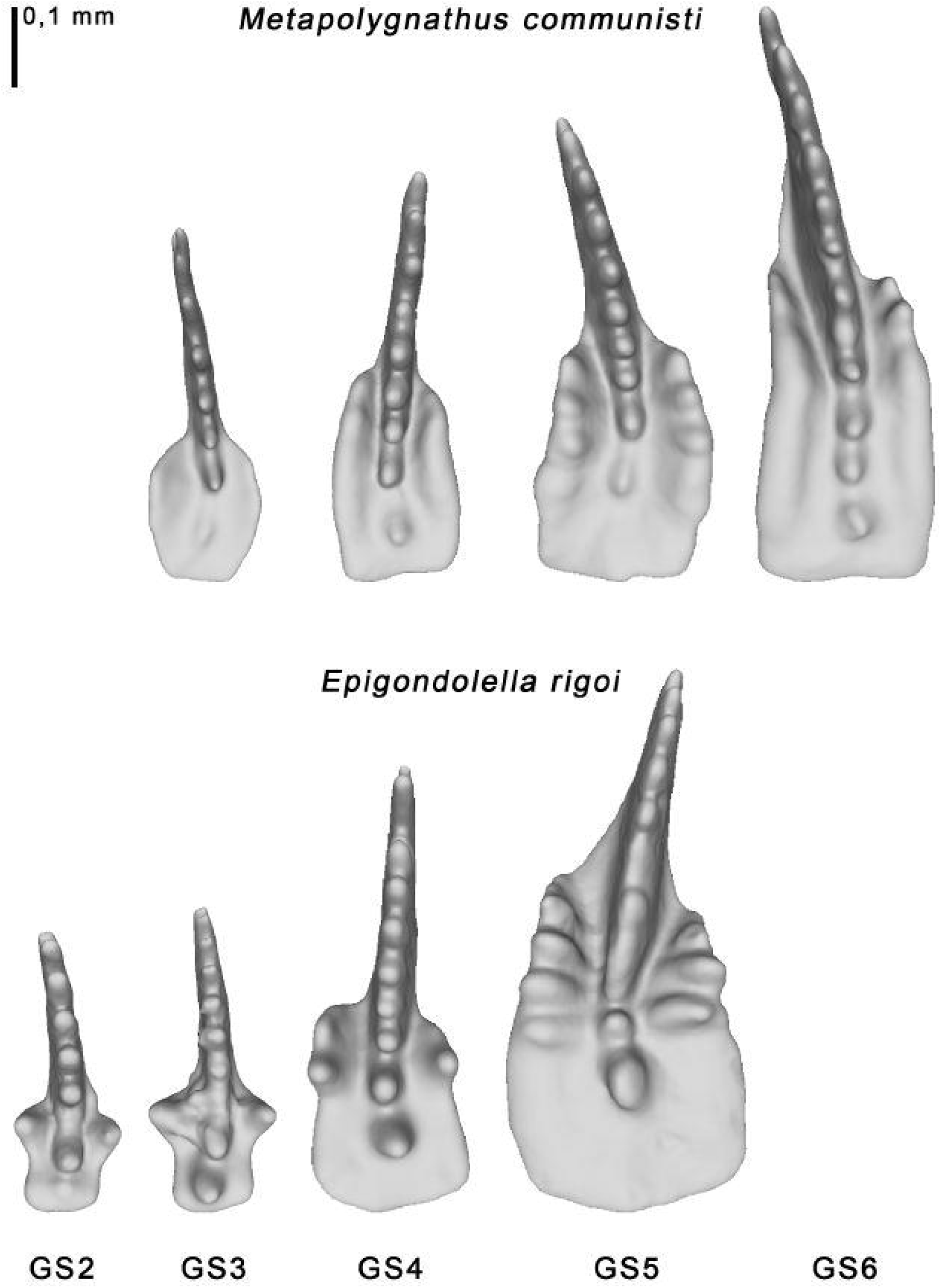
Growth stages of *Metapolygnathus communisti* and *Epigondolella rigoi*. For *M. communisti*: **GS3** – specimen NA37_18; **GS4** – specimen NA37_12; **GS5** – specimen NA37_01; **GS6** – specimen NA37_24. For *E. rigoi*: **GS2** – specimen NA59_20; **GS3** – specimen NA59_19; **GS4** – specimen NA59_01; **GS5** – specimen NA59_05.

#### Growth allometry

The length of the element was used as a proxy for the size of the conodont animal, as was done in previous studies (Purnell 1993, 1994; Zhang et al. 2018; Ginot and Goudemand 2019). The element length, the platform length and the platform area were measured using the 3D software MeshLab 2020.12 (Cignoni et al. 2008). The length of the element was measured from the anteriormost point of the blade in a straight line to the middle of the posterior edge of the element’s platform (Figure 3). As the platform is not equally long on the two sides of the blade, its length was measured in two ways. First, as most elements are curved, the convex side of the platform was measured. In *Metapolygnathus communisti*, the convex size tends to be the longer side of the platform, though not always. In *Epigondolella rigoi* the convex side was almost exclusively the longer side. Alternatively, the longest side of the platform was measured, regardless of curvature (Figure 3, see also Figure S1 in Supplementary Information). In *M. communisti*, in both instances, the platform was measured from the most anterior part of the platform to its posterior end in a line parallel to the imagined symmetrical axis of the platform (Figure 3). In *E. rigoi*, the platform was measured from the geniculation point to the platform’s posterior end. This measure was chosen because the anterior trough margin in this species, though reduced, reaches quite far up the blade, especially in more mature growth stages (see Mazza et al. 2012a for details about the taxonomic characters).

**Figure 3:**
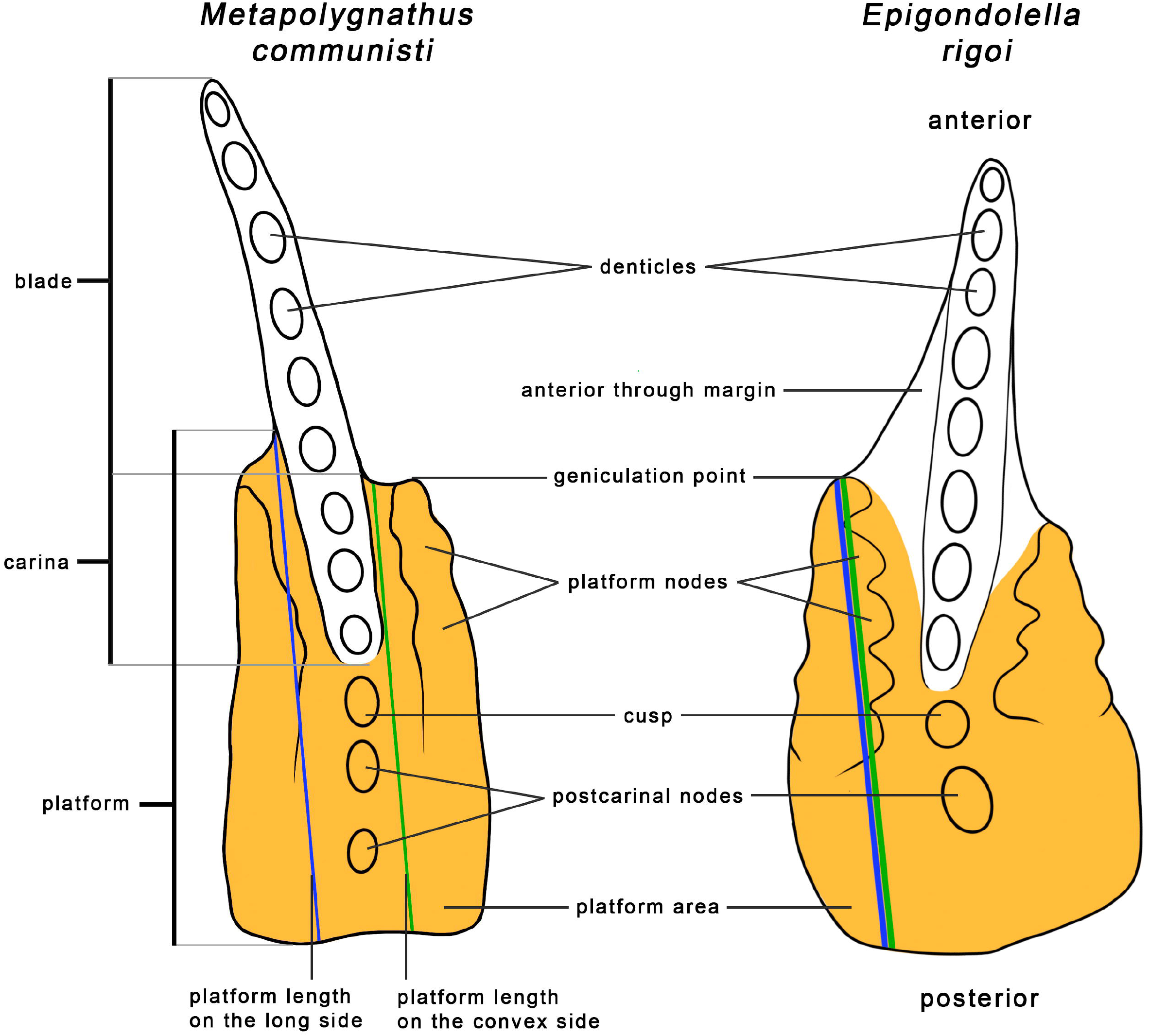
Measurements and morphological characters of P_1_ elements illustrated on *Metapolygnathus communisti* specimen NA37_24 and *Epigondolella rigoi* specimen NA59_05. In most cases the convex side is also the long side. Morphological characters follow Mazza et al. (2012a, b).

The measured area of the platform includes the platform, the cusp, and any postcarinal nodes (Figure 3). In *M. communisti*, specimens of earlier growth stages tend to exhibit only one postcarinal node, already present in GS1 (Mazza and Martínez-Pérez 2015). From GS3 on, a second posterior node may appear (Mazza and Martínez-Pérez 2015). Adult specimens show three or four posterior carinal nodes (Mazza et al. 2012*a*). However, our measurements consistently included only two or three postcarinal nodes in adult specimens. In *E. rigoi*, the cusp is always followed by a single larger postcarinal node (Mazza et al. 2012a). These parts of P_1_ elements were added to the measurements of the platform area, even though they are taxonomically not part of the platform because they likely played a similar part in the processing of food as the platform itself. In *E. rigoi*, the anterior trough margin was not included in the measurements of the platform area (Figure 3). The anterior trough margin is not present in *M. communisti* (Mazza et al. 2012a).

Reduced major axis regression (RMA) was calculated using the R package “smatr” 3.4-8 (Warton et al. 2012; R Core Team 2021) to examine the relationship between the length of the platform and the length of the element, as well as the platform area and the length of the element. All measurements were log-transformed to assess allometric relationships. RMA was chosen as a method because both variables are mutually dependent. Slopes obtained with *E. rigoi* and *M. communisti* data were compared with the “slope.com” function of the “smatr” R package. The same function was used to compare slopes related to the convex side and the longer side of the platform within a species. Slopes between both species and isometry were compared with “slope.test” function of “smart” R package. Isometry was modeled with a slope coefficient of 1 when testing the platform length *vs*. element length; a slope coefficient of 2 when testing the platform area *vs*. element length. In the case slopes do not differ significantly, we compare the intercepts and their confidence intervals showed with the “sma” function. A differentiation in trophic niche could be indeed reflected in different intercepts values when slopes are similar (Lumer et al. 1942; Gould 1979).

Purnell (1994) used a Z-test (Hayami and Matsukuma 1970) to test whether slope coefficients of *Idiognathodus* sp. and *Gnathodus bilineatus* differed significantly from isometric growth. A Z-index higher than 1.96 means that the relationship differs from isometry significantly. We consider this index comparable to the p-values we obtained for *M. communisti* and *E. rigoi* when comparing their slope coefficients with slopes expected under isometry. To compare slope coefficients from this study with those provided by Purnell (1994), 95% confidence intervals (95% CI) of slope coefficients of *Idiognathodus* sp. and *G. bilineatus* were calculated according to the following formula:

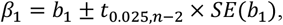

Where *b*_*1*_ is the slope coefficient, *n* is the number of measurements, *t* is the t distribution and *SE* – standard error. Measurements for the platform length of *G. bilineatus* were unavailable (Purnell 1994).

#### Quantitative topographic analysis (DNE)

Dirichlet Normal Energy (DNE) measures a surface’s curvature (Bunn et al. 2011). The DNE of an object is independent of its size and orientation. Its equation is commonly written as follows:

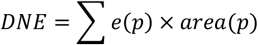

Here, *e(p)* is the Dirichlet Energy Density at a point *p*. The sum of the areas of all points *p* (however small) on a surface is equal to the total area of the surface. Flat planes have DNE values of 0. Therefore, a higher DNE value expresses the elements’ complexity and the average “sharpness” of a surface (Bunn et al. 2011).

We applied the DNE on P_1_ element meshes with the R package “molaR” 5.0 (Pampush et al. 2016). To ensure consistency of DNE calculation, all meshes were simplified to 10 000 faces (Spradley et al. 2017) using Quadric Edge Collapse Decimation in MeshLab. They were then rescaled so that each platform area equaled 0.1 mm^2^. Meshes were then smoothed in Avizo using the Avizo smoothing function with lambda = 0.6 and 25 iterations. Twenty to thirty smoothing iterations, a conservative amount when compared to other approaches (Bunn et al. 2011; Winchester et al. 2014; Spradley et al. 2017), are recommended because they eliminate scanning noise while capturing fine-scale features and avoiding the creation of artificial dimples that can be caused by over smoothing (Spradley et al. 2017). Different numbers of iterations (5, 10, 15, 20, 25, 30, 40, 50, 60, 70, 80, 90, and 100) were tested on a single specimen of *M. communisti* to determine the impact of the number of smoothing iterations (Figure S2). The DNE appears stable from ∼20 to 25 iterations.

The meshes were then manually cut to keep the occluding surface of the platform, the cusp and the postcarinal nodes (Figure 4). Additionally, we cut out the aboral part of the platform because it does not take an active part in food processing. The meshes were finally imported into R Software as binary *ply* files. Individual pieces of meshes created by the smoothing operation were removed to prevent them from affecting the DNE calculation. DNE was calculated with an included boundary exclusion criterion (BoundaryDiscard=“vertex”), as advised by Spradley et al. (2017). The total surface DNE is the mean of all DNE values for individual faces of a surface (Pampush et al. 2016). Because a DNE value is not a traditional measurement, we did not log-transformed the data. We rather fitted the relationship between DNE values and element dimensions (i.e element length, platform length and platform area) with a power law using the “aomisc” R package (Onofri 2020). In addition, Growth stage 5 (GS5) was chosen to compare DNE values between species because this stage comprised enough specimens from both species to allow statistical analysis (Table 1). GS5, representing adults, also allows for interpretations of the diet. The medians of DNE distributions were compared with a Kruskal-Wallis test in R (Hollander and Wolfe 1973).

**Figure 4:**
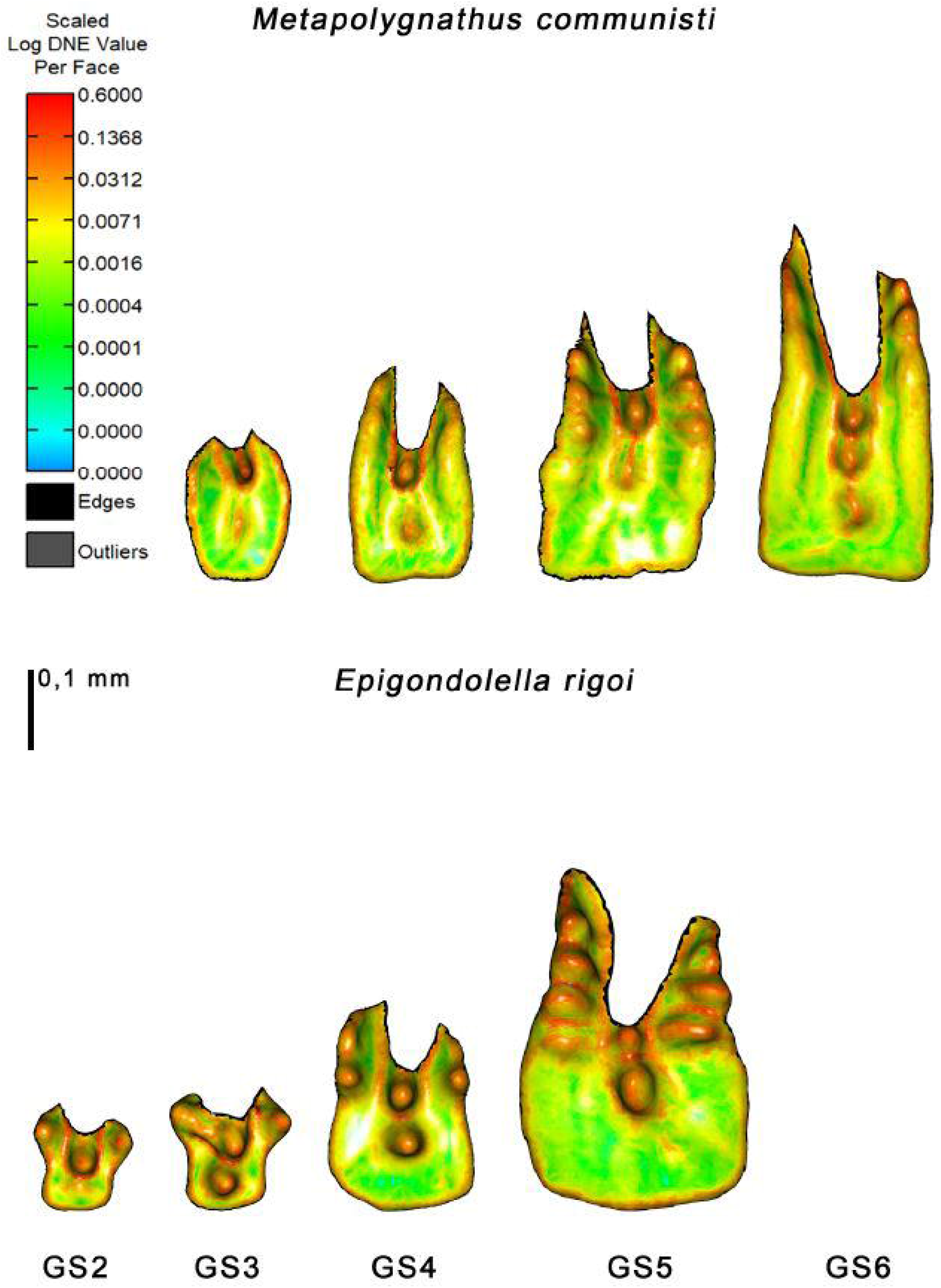
Log-transformed DNE values across growth stages GS3 to GS6 of *Metapolygnathus communisti* and across growth stages GS2 to GS5 of *Epigondolella rigoi*. Names of the specimens are specified in Figure 2.

The R code and data used for investigating allometric patterns and DNE are available on OSF at this address: https://osf.io/283wq/?view_only=6f22274998134eb99cbe43749c6e3e7e (Kelz et al. 2022).

## Results

### Growth allometry

In all examined relationships, high coefficients of determination (R^2^ ≥ 0.89) indicated that linear regression captured the relationships between the variables sufficiently (Table 2). In *Metapolygnathus communisti*, linear regression slopes of the platform length over the P_1_ element length and the platform area over the P_1_ element length showed positive allometry (Figure 5A-B). In both cases, the slopes differed significantly from values corresponding to isometry, i.e. one and two, respectively (Table 2). No differences between the two ways the platform length was measured could be detected (p=0.23; Figure S1A).

**Table 2.** Linear regressions for platform length and platform area over element length for each studied conodont species. Slope coefficients and R^2^ result from the Reduced Major Axis method. The p-value results from the “slope.test” function that compared the coefficients between species and isometry.

| Regression model | Taxon | Slope | Slope CI lower value | Slope CI upper value | R<sup>2</sup> | p-value (h0 = isometric growth) | Intercept | Intercept CI lower value | Intercept CI upper value |
| --- | --- | --- | --- | --- | --- | --- | --- | --- | --- |
| Platform length over element length | *Metapolygnathus communisti* | 1.580 | 1.402 | 1.781 | 0.92 | $1.74 \times 10^{-8}$ | -0.271 | -0.361 | -0.180 |
| | *Epigondolella rigoi* | 1.403 | 1.238 | 1.590 | 0.92 | $1.10 \times 10^{-5}$ | -0.428 | -0.538 | -0.318 |
|  | *Idiognathodus* sp. | 1.089 | 1.011 | 1.167 | 0.98 | 2.345 (Z test) |  |  |  |
| Platform area over element length | *Metapolygnathus communisti* | 2.561 | 2.230 | 2.940 | 0.89 | $1.039 \times 10^{-3}$ | -1.376 | -1.545 | -1.206 |
| | *Epigondolella rigoi* | 2.392 | 2.080 | 2.750 | 0.90 | $1.443 \times 10^{-2}$ | -1.229 | -1.439 | -1.019 |
|  | *Idiognathodus* sp. | 2.149 | 2.002 | 2.296 | 0.98 | 2.060 (Z test) |  |  |  |
|  | *Gnathodus bilineatus* | 2.164 | 2.020 | 2.308 | 0.98 | 2.329 (Z test) |  |  |  |

**Figure 5:**
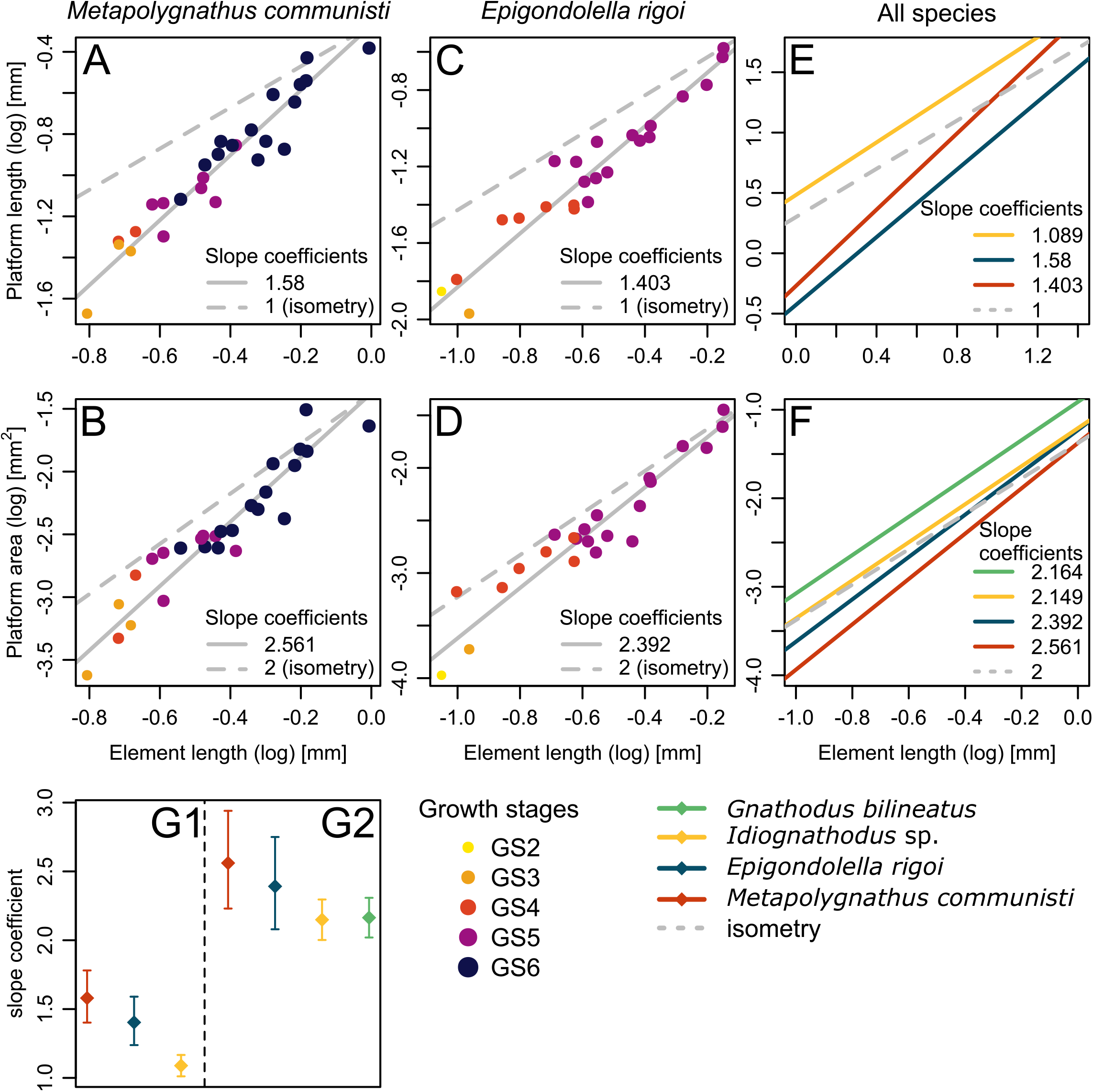
**A-D**. Allometric growth of P_1_ platform length and P_1_ platform area for *Metapolygnathus communisti* (n=27) and *Epigondolella rigoi* (n=23); each dot represents an element and the colours are related to the growth stage. **A**. Platform length over element length for *M. communisti*; convex platform length only. **B**. Platform area over element length for *M. communisti*. **C**. Platform length over element length for *E. rigoi*; convex platform length only. **D**. Platform area over element length for *E. rigoi*. **E-F**. Comparison of the allometric growth between *Gnathodus bilineatus* (Purnell 1994), *Idiognathodus* sp. (Purnell 1993, 1994), *E. rigoi* and *M. communisti*. **E**. Platform length over element length. Platform length in *M. communisti* and *E. rigoi* was measured on the convex side. No data were available for *G. bilineatus* (Purnell 1994). **F**. Platform area over element length. **G**. Comparison of slope coefficients between species; G1, Platform length over element length; G2, platform area over element length.

In *Epigondolella rigoi*, both regression slopes (platform length over element length and platform area over element length) indicated positive allometry and were significantly higher than predicted under the null hypothesis of isometric growth (Table 2; Figure 5 Morphological characters follow Mazza et al. (2012a, b).

Figure 4. : Log-transformed DNE values across growth stages GS3 to GS6 of *Metapolygnathus communisti* and across growth stages GS2 to GS5 of *Epigondolella rigoi*. Names of the specimens are specified in Figure 2.

Figure 5C-D). No significant difference between the two ways the platform length was measured could be detected (p=0.83; Figure S1B).

Slope coefficients of the platform length over element length did not differ significantly between *M. communisti* and *E. rigoi* (p=0.166), but in both cases, they were significantly higher than that of *Idiognathodus* sp. as their confidence intervals did not overlap (Table 2, Figure 5E, G). Platform areas in all four species showed positive allometry over element length (slope coefficients between 2.149 and 2.561) but the four 95% confident intervals overlapped, and no significant difference was detected between *M. communisti* and *E. rigoi* (p=0.479). No differences in intercepts could be detected for any of the examined relationships between *E. rigoi* and *M. communisti* (Table 2).

### Dental topography (DNE)

In *M. communisti*, DNE values ranged between 99.93 and 279.84; in *E. rigoi*, between 117.59 and 353.71 (Figure 6). Specimens classified as GS5 of *E. rigoi* showed higher DNE values than GS5 specimens of *M. communisti* (Kruskal-Wallis test, p=0.015; Figure 6A).

**Figure 6:**
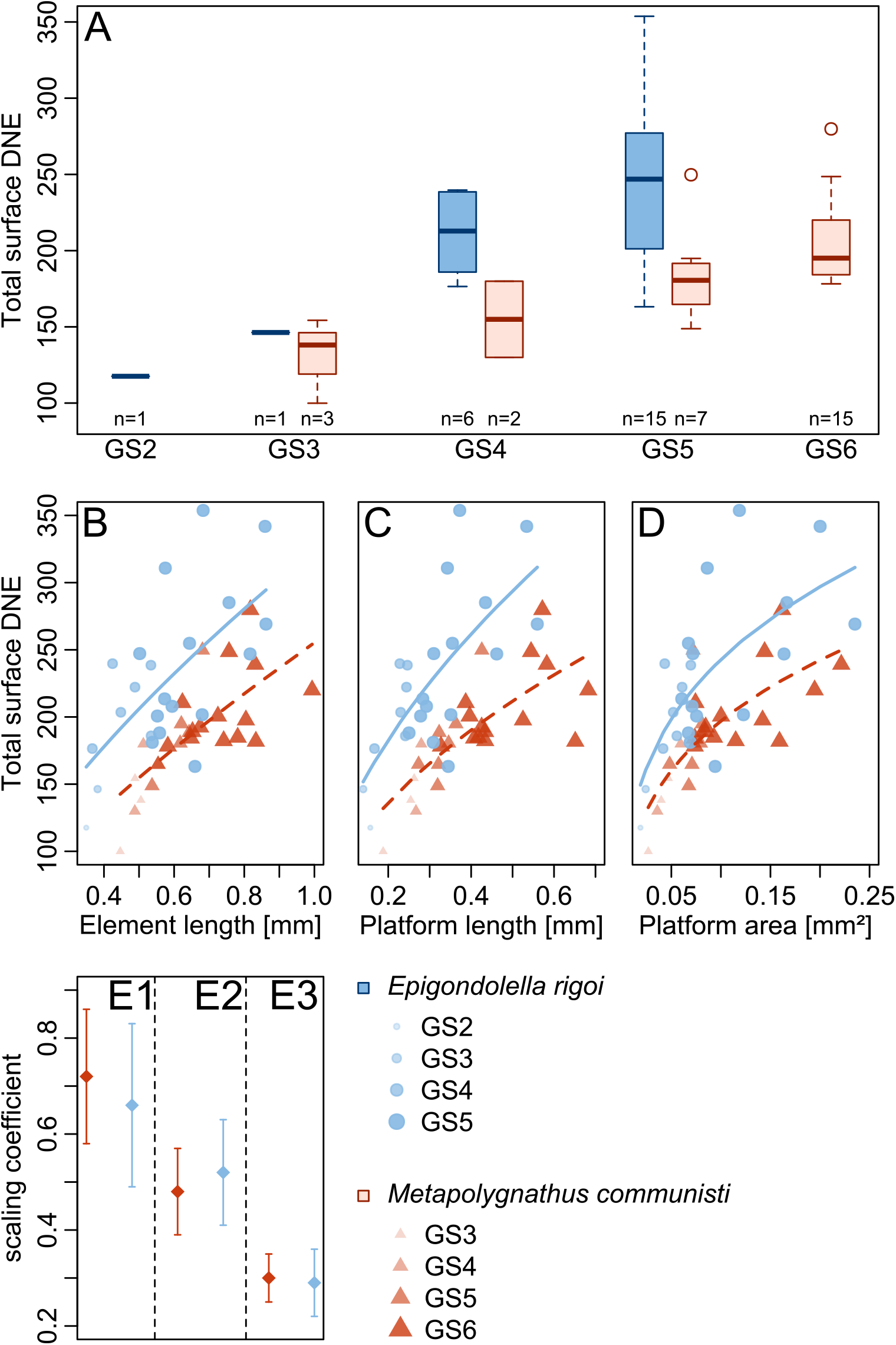
**A**. Distribution of DNE values across growth stages in *E. rigoi* and *M. communisti*. **B**. DNE values over element length with fitting to the allometric equation *y = aX*^*b*^. **C**. DNE values over platform length with fitting to the allometric equation *y = aX*^*b*^. **D**. DNE values over platform area with fitting to the allometric equation *y = aX*^*b*^. **E**. Comparison of scaling exponents between species; E1, DNE values over element length; E2, DNE values over platform length; E3, DNE values over platform area.

The relationship between DNE values and P_1_ dimensions show a modest fit to the power law (0.42 ≥ R^2^ ≥ 0.59; Table 3, Figure 6B-D).Figure 6 Between *E. rigoi* and *M. communisti*, the scaling exponents (“b” in Table 3) are respectively 0.66 and 0.72 in DNE *vs*. element length; 0.52 and 0.48 in DNE *vs*. platform length; and 0.29 and 0.30 in DNE *vs*. platform area (Table 3). No difference in exponents is detected for any of the examined relationships (Figure 6E).

**Table 3.** Parameters of the allometric equation *y = aX*^*b*^ fitted to DNE values over element dimensions for each studied conodont species. All values have been rounded to the second significant digit.

| Regression model | Species | a | SE(a) | b | SE(b) | R<sup>2</sup> |
| --- | --- | --- | --- | --- | --- | --- |
| DNE over element length | *Metapolygnathus communisti* | 255.25 | 15.54 | 0.72 | 0.14 | 0.54 |
|  | *Epigondolella rigoi* | 324.71 | 30.92 | 0.66 | 0.17 | 0.42 |
| DNE over platform length | *Metapolygnathus communisti* | 296.15 | 23.56 | 0.48 | 0.09 | 0.59 |
|  | *Epigondolella rigoi* | 420.53 | 55.22 | 0.52 | 0.11 | 0.51 |
| DNE over platform area | *Metapolygnathus communisti* | 392.06 | 49.79 | 0.30 | 0.05 | 0.59 |
|  | *Epigondolella rigoi* | 474.07 | 78.79 | 0.29 | 0.07 | 0.48 |

## Discussion

The null hypothesis of an isometric growth of P_1_ elements in *M. communisti* and *E. rigoi* could be rejected based on positive growth allometry of the platform length and platform area over element length. These results support previous findings of positive growth allometry in these organs in much older, Carboniferous, ozarkodinid taxa *G. bilineatus* and *Idiognathodus* sp. (Purnell 1993, 1994). Under the assumption that positive allometry reflects the organism’s energy demand growing at a higher pace than the surface of a (molar) tooth (Gould 1966, 1975), these results are consistent with the interpretation of P_1_ elements as organs used for mechanical slicing and grinding of food, as previously proposed based on microwear (Purnell 1995; Martínez-Pérez et al. 2014b), enamel-like ultrastructure of conodont lamellar crown tissue (Donoghue 2001b), and finite element analysis (Jones et al. 2012a). However, the assumptions of the original studies by Purnell (1993, 1994) need to be revisited in the context of the studies which had been made in the field of metabolic ecology over the last 20 years. The premise of the argument based on tooth positive allometry was that an animal’s metabolic rate, and thus energy requirements, increase as its volume to the power of 0.75. This scaling exponent has been predicted by several theories (Kleiber 1932; Feldman and McMahon 1983; West et al. 1997; Savage et al. 2004) but an exponent of 0.67 has been equally reported in mammals and derived from theoretical considerations (Heusner 1982; Dodds et al. 2001; White and Seymour 2003), so the same exponent that in assumption 3 developed in the Introduction. This disagreement has important implications for the reasoning in the studies by Purnell (1993, 1994): if the metabolic rate in conodonts scaled as body mass to the power of 0.67 and not 0.75, then an isometric relationship between the food acquisition surface and the animal’s dimensions might well be expected. In such case, it would be impossible to distinguish whether positive allometry reflects differences in scaling between energy demand and food-processing tooth surface or whether the two variables are not related. The latter possibility would preclude the use of growth allometry in inferring trophic ecology in fossil organisms. We attempt, therefore, to extrapolate the relationship in conodonts by comparing with related organisms. The average scaling exponent in fish has been reported as 0.79 (Clarke and Johnston 1999)or even 0.99 (Glazier 2009). In ectoterms, exponents higher than 0.75 have been reported across all groups compiled by Glazier (2007). Based on analogy, a scaling exponent of metabolic rate over body mass or volume can be in conodonts expected to be higher than 0.67. It should be, however, kept in mind that this is a strong and untestable assumption, as the scaling exponents vary extremely between animals and even within taxa, depending on their interactions with other organisms (Bokma 2004; Glazier 2009; Sieg et al. 2009). The scaling exponents of DNE over element length lie between 0.66 and 0.72, thus closer to the 0.67 exponent than to 0.75. This might serve as an indication that in conodonts, energy demand grew in a more mammal-like fashion (closer to M^0.67^) than in a fish or reptile fashion (M^>0.67^), in line with the idea of conodont P_1_ elements functioning analogously to molars (Donoghue and Purnell 1999*b*). But scaling of DNE with tooth size has only been recently identified apes (Berthaume and Schroer 2017) and the interpretation of the relationships between this metric, body size and metabolism are little known, therefore the scaling observed here requires further comparisons with extant organisms. Here we tentatively uphold the reasoning and conclusions by Purnell (1993, 1994) and extend them to the new taxa examined here: if the metabolic rate in conodonts scaled as M^>0.67^, then the positive allometry of conodont element dimensions supports the grasping tooth model in all four taxa and the test may be used in other conodont element morphologies.

### Functional differences between conodont species

The allometric relationship between platform area and element length did not differ significantly between the two Triassic species examined here and Carboniferous taxa *Idiognathodus* sp. and *G. bilineatus* (Figure 5G2). However, there is a significant difference between Triassic species and *Idiognathodus* sp. when considering the growth of the platform length (Figure 5G1). Perhaps this difference in slope coefficient resulted from the methodological differences in how platform measurements were taken here and in Purnell (1993, 1994). Measurements based on pictures often suffer from distortions from projecting a 3D structure onto a plane, where differences in levelling the photographed specimens might affect the results (Mullin and Taylor 2002; Collins and Gazley 2017). Proper 3D measurements of P_1_ platform length and area using our methodological protocol should be investigated in *Idiognathodus* sp. and *G. bilineatus* to verify any difference in allometric slope coefficients.However, as platform length showed a stronger positive allometry in Late Triassic taxa than in *Idiognathodus* sp., it suggests that the increase of functional surface was primarily achieved by platform elongation, rather than growth in width.

Our hypothesis that *M. communisti* and *E. rigoi* differed in the growth allometry of their P_1_ elements could not be rejected (Table 2). Similar growth allometry might indicate similar energy demand, as well as similar efficiency in food acquisition and assimilation.

### Conodont diet inferred by platform curviness

Previous investigations showed that dental topography methods could be applied to non-homologous dental tools to track dietary differences between distantly related clades (Stockey et al. 2021). Conodont elements in this study showed similar curviness to complex mammal teeth. This similarity does not mean that direct dietary associations can be made between conodonts and mammals, but dental topography allows comparisons between taxa and ontogenetic stages and helps in constraining conodont ecology (Purnell and Evans 2009). In *M. communisti* and *E. rigoi*, DNE values differed significantly at the adult growth stage (i.e., GS5). We tested the differences in DNE values only at GS5 because ontogenetically younger growth stages were represented by fewer specimens in both species (Table 1), which made observations of these stages less conclusive. Differences in GS5 DNE values between *M. communisti* and *E. rigoi* allow rejecting the hypothesis that adult specimens of both species shared the same diet. More DNE analyses on conodonts are needed to understand the scope of DNE values in this group and to confidently suggest that a discrepancy between DNE values of different species reflects different dietary niches. Indeed, DNE values of GS5 specimens of *M. communisti* are similar to those reported for folivores or omnivores; insectivores or folivores in the case of *E. rigoi*, but these diets based on DNE values stem from studies on primates (Bunn et al. 2011; Winchester et al. 2014). Moreover, conodont elements are more than ten times smaller than primate teeth, complicating dietary comparison as conodonts could not eat the same objects. Though primate dietary classifications do not apply to conodonts, they may offer a general reference point for the methods of breaking down different food types. Insectivores rely on sharp cusps to apply maximal force to a small surface area to pierce hard insect chitin, and folivores also use steeply sloped cusps to shear tough cellulose-rich leaves (Lucas 1979; Strait 1997). Therefore, it is possible that conodont element platforms evolved to break down food types with similar mechanical properties. For instance, conodonts may have punctured arthropod larvae (Dzik 2021), a diet consistent with the DNE values observed in *M. communisti*.

### Current limits on DNE and advices for further investigations

The definition of the platform for DNE measurement is worth discussing because it is somewhat subjective. In this work, we decided to include the cusp and all postcarinal nodes because, in many cases, the cusp marked a notable transition between sharper denticles of the blade and flatter nodes on the platform. In *M. communisti*, the number of postcarinal nodes increases through growth (Mazza et al. 2012a; Mazza and Martínez-Pérez 2015), impacting DNE values through the ontogeny of this species. In *E. rigoi*, the number of posterior nodes stays the same (Mazza et al. 2012a), so variations of DNE values are more related to the growth of prior postcarinal nodes or the addition of new nodes on the edge of the platform. This difference in node location on the platform might affect the way conodonts broke down food but it should be further investigated by assessing occlusal kinematics of *M. communisti* and *E. rigoi* P_1_ elements, which is currently impossible because no clusters (i.e., a conodont apparatus with elements found in connection) are currently known for these species.

There are several challenges when applying DNE to conodonts. As DNE is a comparatively new tool, reference values and understanding of variability (e.g. intraspecific, ontogenetic and taphonomic) are limited. So far, DNE research has focused on mammals, whose tooth function largely relies on jaws acting as levers. The different mechanics of feeding in jawed organisms necessitates that the comparisons drawn here between conodont elements and primate molars must be viewed as extremely hypothetical. Marine environment, evolutionary distance and the lack of jaws in conodonts make tooth function likely not completely analogous between the two. Even more importantly, conodonts are unique among vertebrates in repairing their teeth by periodic apposition of new growth layers on top of the ones previously used for food processing (Shirley et al. 2018). This mechanism is both used to repair damage and restore sharpness (Donoghue, 1998), as well as to generate topographic complexity (Müller and Nogami 1971).

Thus, DNE values in conodont elements will increase as the elements grow and so the animal gets older. This is not the case in mammals due to their lack of tooth repairing ability: it has been showed with mice’s molars that DNE values decrease with the age of the animal because of teeth abrasion (Savriama et al. 2022). That is why typical applications of DNE on mammals do not consider ontogenetic development, authors rather investigate DNE variation between young adults from different species, or between different teeth within one specimen. For example, Pérez-Ramos et al. (2020) compared DNE values between premolars and molars in cave bears, and they suggested that “increasing upper tooth surface areas also increase the values of both topographic variables, and probably, also their chewing efficiency”.

A consistent protocol of mesh preparation before DNE calculation is also needed to get a better reproducibility of the method. For example, the scan resolution and the number of smoothing iterations vary in current literature, which can impact DNE values (Spradley et al. 2017; Assemat et al. 2022). This variation in processing is important because comparisons are most conclusive when drawn between data with similar preparation. Furthermore, the scan resolution depends on the size of the studied object. We cannot expect a 1 μm resolution (as for conodont P_1_ elements from the Pizzo Mondello (Guenser et al. 2019)) for mammal teeth of several millimetres in length. However, we can set a standardized scan resolution for conodont elements of 1 μm (even lower would be better), which would allow investigating ontogenetic patterns by including P_1_ elements of less than 400 μm in length (i.e., juveniles). About the post-scanning preparation, we second Spradley et al.’s (2017) recommendations: a conservative number of smoothing iterations (20-30) using non-Laplace-based smoothing operators, such as that implemented in Avizo, and mesh simplification to a fixed number of faces.

## Conclusions

We used 3D meshes to test the null hypothesis that Late Triassic conodont P_1_ elements grew isometrically, thus revisiting Purnell’s (1993, 1994) test of conodont element function. We tested this hypothesis against the alternative that the elements showed positive allometry. Positive allometry had been proposed to be a test of the grasping-tooth hypothesis (Aldridge et al. 1987; Purnell and von Bitter 1992). However, the test requires knowledge of the scaling exponent of the metabolic rate to body mass (volume). We followed Purnell’s protocol (1993, 1994), analyzing the growth allometry of the platform length and area *vs*. total length of the element, originally performed on 2D projections of conodont elements. Platform length and platform area showed positive allometry relative to element length, allowing us to reject the null hypothesis. If metabolic rate in conodonts scaled with body mass similarly to that in fish and ectoterms, the test holds and our results support the grasping-tooth hypothesis. Slope coefficients nor intercepts did not differ between *M. communisti* and *E. rigoi*. However, for platform length over element length, slope coefficients were higher than those reported for Carboniferous conodonts *Idiognathodus* sp. (Purnell 1994), which may suggest that the energy demand (and thus metabolic rate) increased faster in this species as the organism grew. A more precise measurement of surface area in 3D models compared to 2D projections used in *Idiognathodus* sp. did not result in a significantly different relationship. In contrast, length-rather than area-based measurements allowed detecting significant differences between Late Triassic taxa and *Idiognathodus* sp.

We added dental topographic analysis of the platforms using DNE to test the hypothesis that co-occurring species *M. communisti* and *E. rigoi* shared the same food base. *E. rigoi* showed significantly higher DNE values than *M. communisti* when comparing adult growth stages.Although DNE is a size-independent measure, its values increased with the size of adult specimens in both Triassic taxa. This increase is interpreted here as the reflection of energy demand growing faster than the area of the element and thus potentially reflecting the scaling of the metabolic rate consistent with the assumptions of the studies by Purnell (1993, 1994). Scaling exponents of DNE over element length in an allometric model were closer to those reported for mammals than for fish and other ectoterms. As these exponents did not differ substantially between *M. communisti* and *E. rigoi*, the differences in DNE are not likely to arise from differing metabolic rates and can be attributed to different food bases.

Our study revisited one of the main arguments used in interpreting the function of conodont elements: their positive growth allometry. We argue that a key assumption of this interpretation, the scaling of metabolic rate as mass (volume) to the power of 0.75, cannot be made for conodonts automatically, as there is substantial variation between known animals. We also show that co-occurring taxa differed in their diets, as reported previously in Silurian conodont communities (Terrill et al. 2022), which supports trophic diversification as an important driver of the remarkable disparity of their elements.

## Supporting information

Kelz et al_supp mat

## Acknowledgements

We thank Bryan Shirley for support in using Aviso and Wyatt Petryshen for advice on cleaning and saving meshes. We also thank Michele Mazza for their advice and field sampling. We finally thank Nicolas Goudemand for supporting the project (French ANR grant, ACHN project EvoDevOdonto). PG was supported by Visiting Scholarship awarded by Friedrich-

Alexander-Universität Erlangen-Nürnberg. EJ was supported by Deutsche Forschungsgemeinschaft (project no JA 2718/3-1). MR was supported by DOR2054230/20 by University of Padova. We are grateful to Viktor Karádi, Nicolas Campione and Paleobiology Editor James Crampton for constructive comments, which improved the manuscript, and to Craig White for sharing data on the scaling of metabolic rates in various taxa.

## Table captions

Table 1. Numbers of conodont P1 element specimens by growth stages used for the study. Abbreviations: GS – Growth stage.

Table 2. Linear regressions for platform length and platform area over element length for *Metapolygnathus communisti, Epigondolella rigoi, Idiognathodus* sp. and *Gnathodus bilineatus*. Data for *Idiognathodus* sp. and *G. bilineatus* were extracted from Purnell (1994) except for the 95% confidence intervals (CI). Slope coefficients and R^2^ result from Reduced Major Axis method. 95% CI values resulting from “sma” function calculation for *M. communisti* and *E. rigoi*; from calculation for *Idiognathodus* sp. and *G. bilineatus*. The p-value results from the “slope.test” function that compared the coefficients between species and isometry for *M. communisti* and *E. rigoi*; from Z test (Hayami and Matsukuma 1970) for *Idiognathodus* sp. and *G. bilineatus*.

Table 3. Parameters of the allometric equation *y = aX*^*b*^ fitted to DNE values over element dimensions for each studied conodont species. All values have been rounded to the second significant digit.

## Figure captions

