## Supplementary material for "Growth allometry and dental topography in Upper Triassic conodonts support trophic differentiation and molar-like element function": Kelz et al_supp mat

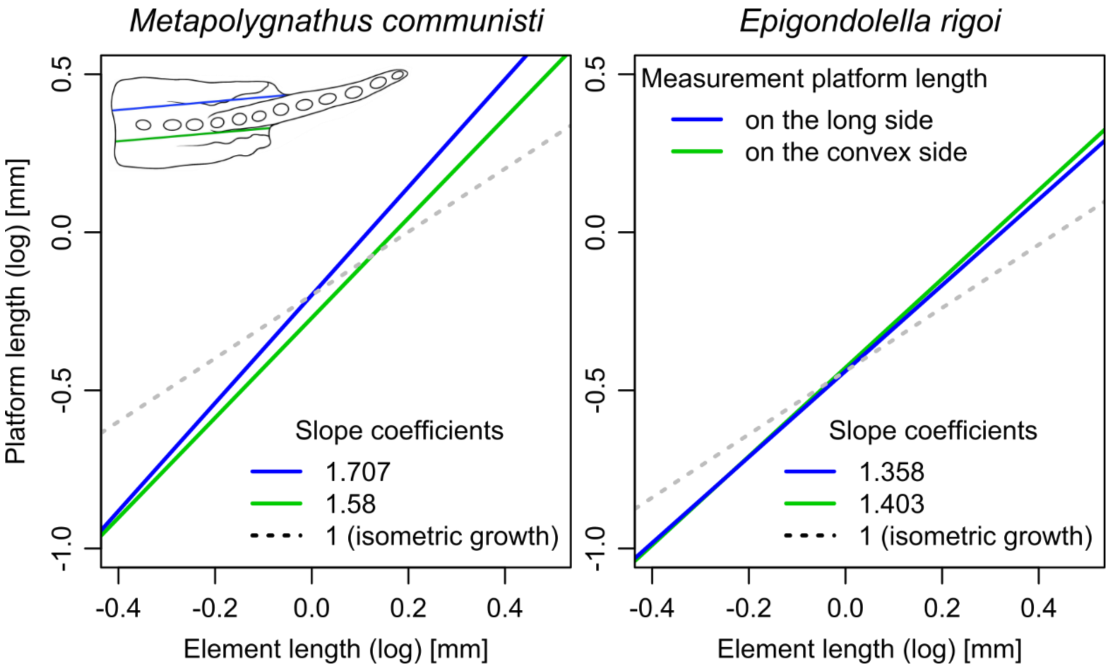


FigS1: Comparison of long and convex platform length on allometric growth of *M. communisti* (n=27) and *E. rigoi* (n=23).

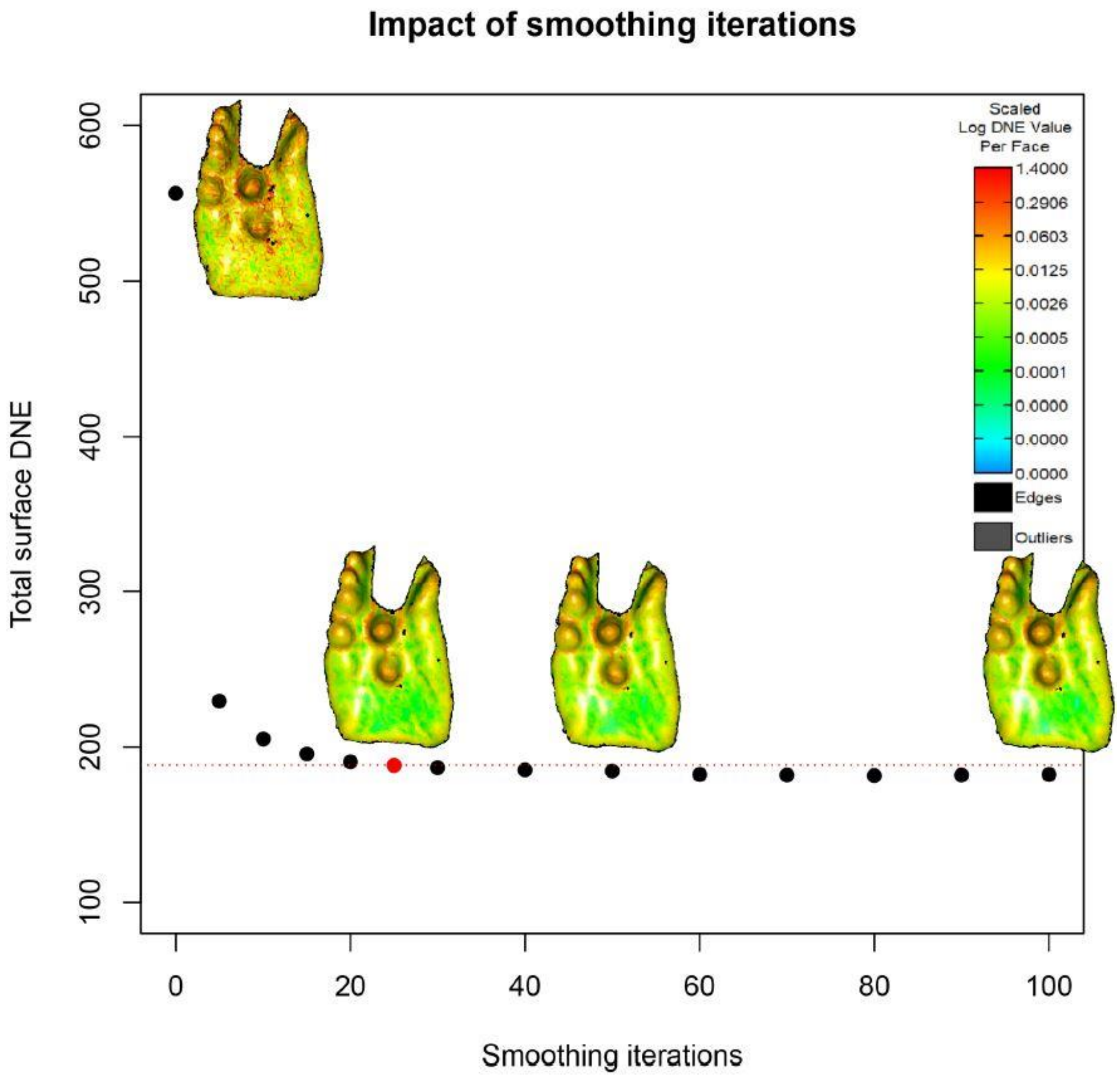


FigS2: The impact of different numbers of smoothing iterations on DNE values measured for the platform of the P1 element specimen NA37_02 (*M. communisti*). The red dot (or grey in black and white printed version) indicates 25 smoothing iterations. For 0, 25, 50 and 100 iterations, a snapshot of a map of log-transformed DNE values applied on a platform of a conodont element (specimen NA37_02) is represented to visualise the impact of elements smoothing on DNE values.
